# Weakest link dynamics predict apparent antibiotic interactions in a model cross-feeding community

**DOI:** 10.1101/2020.03.10.986695

**Authors:** Elizabeth M. Adamowicz, William R. Harcombe

## Abstract

With the growing global threat of antimicrobial resistance, novel strategies are required for combatting resistant pathogens. Combination therapy, wherein multiple drugs are used to treat an infection, has proven highly successful in the treatment of cancer and HIV. However, this practice has proven challenging for the treatment of bacterial infections due to difficulties in selecting the correct combinations and dosages. An additional challenge in infection treatment is the polymicrobial nature of many infections, which may respond to antibiotics differently than a monoculture pathogen. This study tests whether patterns of antibiotic interactions (synergy, antagonism, or independence/additivity) in monoculture can be used to predict antibiotic interactions in an obligate cross-feeding co-culture. Using our previously described weakest link hypothesis, we hypothesized antibiotic interactions in co-culture based on the interactions we observed in monoculture. We then compared our predictions to observed antibiotic interactions in co-culture. We tested the interactions between ten previously identified antibiotic combinations using checkerboard assays. Although our antibiotic combinations interacted differently than predicted in our monocultures, our monoculture results were generally sufficient to predict co-culture patterns based solely on the weakest link hypothesis. These results suggest that combination therapy for cross-feeding multispecies infections may be successfully designed based on antibiotic interaction patterns for their component species.

## Introduction

Antibiotic resistance is a growing global threat. It is estimated that, by 2050, 10 million deaths per year worldwide will be attributable to antibiotic-resistant infections (1). Many previously treatable infections, such as tuberculosis (2), urinary tract infections (3), and even *Staphylococcus-*mediated skin infections (4) now require higher doses of more powerful antibiotics. More concerning is that the patients most at risk for multidrug resistant infections are those with complex medical histories and increased risk of side effects (5). This arms race against pathogens by clinicians is proving a losing battle, as resistance is acquired rapidly, and the development of novel antimicrobials is limited (6, 7). The demand for novel treatment strategies is, therefore, an ever–increasing issue.

One treatment strategy that has proven particularly successful is the use of drug combinations. The best example of this is perhaps for antivirals against HIV, where the advent of highly active antiretroviral therapy (HAART) dramatically improved the longevity and quality of life for HIV patients (8). The theory behind this treatment is based on simple probability — even in a highly mutable and therefore rapid resistance-acquiring virus such as HIV, it is much less likely that a viral population will acquire resistance to multiple antivirals than a single one, assuming an independent mutation is required for resistance to each drug (8, 9). This approach, of using multiple drugs to target multiple essential targets, has also been used in cancer chemotherapy to manage drug-resistant and genetically heterogeneous tumors (10, 11). In cases of bacterial infections, multidrug therapy has been adopted in only a few specific infections, such as treatment for drug sensitive tuberculosis (2). However, clinical trials of combination therapy in the treatment of bacterial infections in patients have been limited. Choosing the correct drug combination is difficult (12, 13), and efficacy has been mixed (14, 15). A greater understanding of the mechanisms driving effective combination therapy are therefore required for successful clinical implementation.

The success of combination therapy is affected by interactions between drugs, wherein the activity and effectiveness of one drug is impacted by the presence or absence of another (16). There are several mechanisms by which antibiotics may synergize (work more effectively or at lower doses together than separately) or antagonize (work less effectively or at higher doses together than separately). While the precise nature of these interactions depends on the drugs and the bacterial species being targeted, some general mechanisms have been described for different classes of antibiotics (17). Synergistic interactions tend to occur when one drug facilitates cellular entry (18–20) or increased efficacy (21) of another, or when the drugs target similar cellular processes (22, 23). Conversely, antagonism may occur when one antibiotic induces tolerance or resistance to another (17, 24, 25), or when one drug corrects for the physiological disruptions caused by another (26). These are general trends only, however, and many species– and drug – specific exceptions apply, making it challenging to predict drug interactions *a priori* in new systems.

Another increasingly appreciated feature of bacterial infections is their polymicrobial nature. Numerous clinically relevant infections are now known to involve multiple species, consisting of a single pathogen and various commensal partners, or several co-infecting pathogens (27, 28). Polymicrobial infections have been observed to have worse clinical outcomes in some cases (29–31), though these results are mixed (32, 33). The metabolic interactions (both positive and negative) among these species have been demonstrated to impact antibiotic response (34). One such positive interaction is cross-feeding, wherein one species produces an essential metabolite for another; this also occurs in infection contexts (35). For example, in a cystic fibrosis model where the pathogen *Pseudomonas aeruginosa* depends on the mucin degradation products supplied by a community of anaerobic commensals, antibiotics specifically targeting the anaerobes decreased *P. aeruginosa* abundance despite its intrinsic resistance to the antibiotic (36). Treatment regimens might, therefore, be more effective if metabolic interactions among species are taken into account; however, little research has been done on how cross-feeding might impact combination therapy.

To this end, we aimed to test whether cross-feeding interactions in a model bacterial community might influence antibiotic interactions. We selected ten combinations of six antibiotics based on the work of Yeh et al. (16); this study quantitatively tested the pairwise interactions between 21 different antibiotics which altered *E. coli* growth rate. Three of the combinations we selected from this study were predicted to synergize (greater antibiotic efficacy in combination than alone); three were predicted to antagonize (lower antibiotic efficacy in combination than alone), and four to interact additively or independently in *E. coli* monoculture. Our model system, consisting of an *E. coli* methionine auxotroph strain that produces acetate from lactose, and an *S. enterica* that produces methionine, has been previously described (36–38).We first tested each of these combinations for their drug interactions in *E. coli* and *S. enterica* monoculture, and used fractional inhibitory concentration indices (FICIs) to identify any drug interactions. We then used our “weakest link” hypothesis to predict the growth patterns of the co-culture and the subsequent antibiotic interactions. Briefly, the weakest link hypothesis states that the “weakest link” species in an obligate cross-feeding community will define the tolerance (i.e. the ability to grow at high antibiotic concentrations) of the entire community. In this study, we found that only three antibiotic combinations showed non-additive interactions; however, weakest link dynamics successfully predicted co-culture growth and antibiotic interactions in these cases. While more antibiotic combinations need to be explored, these results suggest that the responses of individual community members to combination therapy might be sufficient to predict the antibiotic interactions in the larger microbial community.

## Results

Based on previous results in *E. coli* (16), we tested ten combinations of six antibiotics for synergy or antagonism in *E. coli* and *S. enterica* monocultures (**Table 1**). The mechanism of action for each of these antibiotics can be found in **Supplementary table S1.** Each combination was tested in triplicate and minimum inhibitory concentrations (MICs), fractional inhibitory concentrations (FICs) and fractional inhibitory concentration indices (FICIs) were obtained after 48 hours of growth at 30°C (**Figure 1**). To avoid over– or under– interpretation of the antibiotic interactions, we used the median FICI value for each plate and the mean value from each of the three replicate plates for each antibiotic combination.

**Figure 1.**
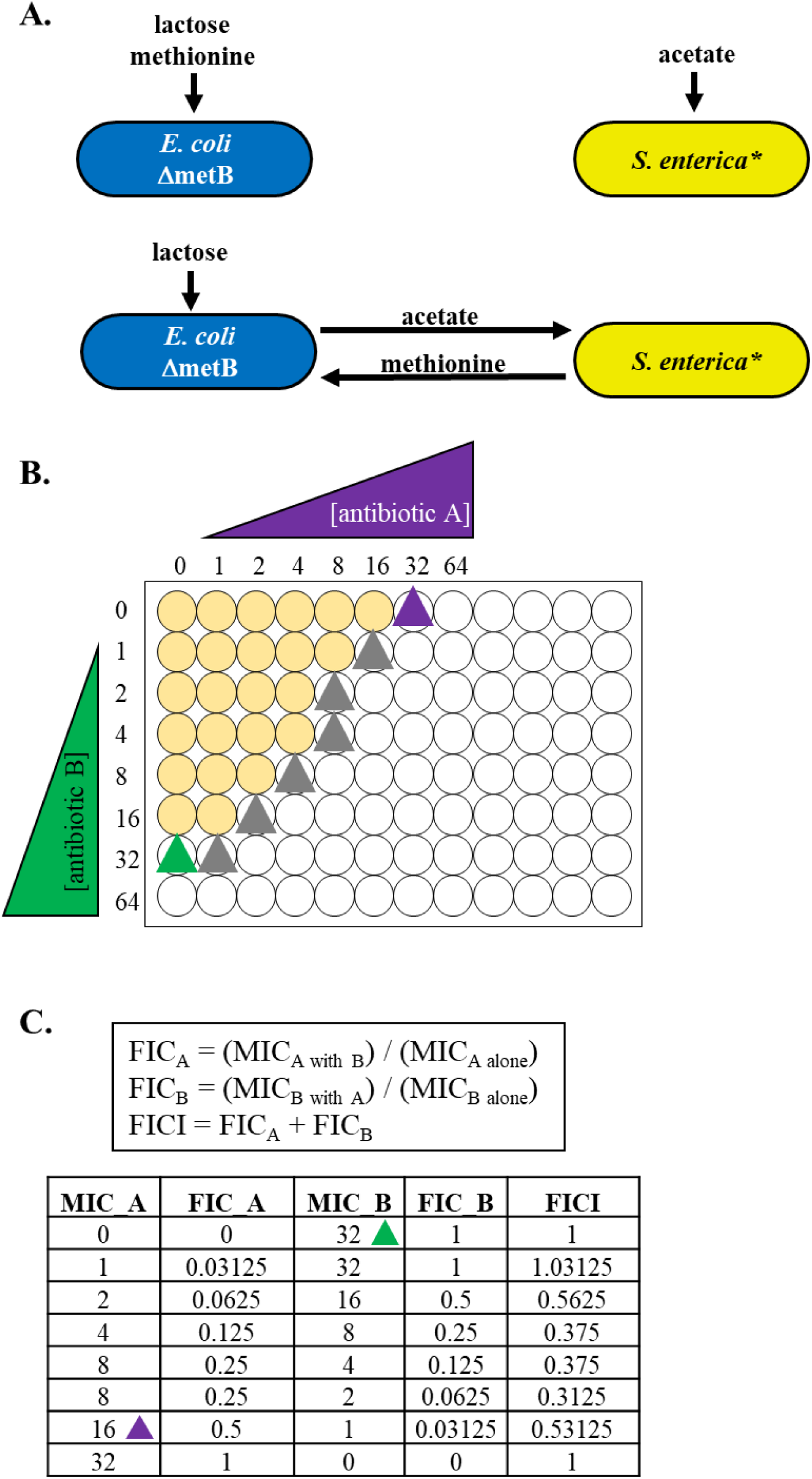
Antibiotic interaction experimental setup and hypotheses. **A.** The two-species obligate cross-feeding system. When lactose is supplied, *E. coli* uses it to produce acetate for *S. enterica*, which produces methionine for *E. coli*. Each species can be grown in co-culture or monoculture, depending on the metabolites supplied. **B.** Setup for checkerboard assays. Seven antibiotic concentrations plus one antibiotic–free well were developed for each antibiotic/ species combination, with the MIC approximately in the middle of the gradient. Mid-log phase cells were inoculated into plates containing species-specific growth medium and antibiotic at twofold dilutions. Cells were allowed to grow for 48 hours at 30°C with shaking, and a Tecan plate reader was used to measure growth at OD600. Growth was defined as an OD600 above 10% of the maximum OD600 obtained on each plate. Three replicates of each antibiotic/ culture condition were obtained. **C.** Table of calculations for fractional inhibitory concentrations and formulae used.

**Table 1.** Antibiotic combinations used in the study and their predicted interactions in *E. coli* based on Yeh et al. 2006.

| Synergy | Antagonism | Additive |
| --- | --- | --- |
| Nalidixic acid and streptomycin | Nalidixic acid and spectinomycin | Nalidixic acid and ciprofloxacin |
| Nalidixic acid and bleomycin | Nalidixic acid and doxycycline | Ciprofloxacin and bleomycin |
| Streptomycin and ciprofloxacin | Spectinomycin and streptomycin | Streptomycin and doxycycline |
|  |  | Spectinomycin and doxycycline |

Previous work from our lab has shown that co-culture growth in the presence of antibiotics is dependent on weakest link dynamics (36). This hypothesis predicts that the MIC of an obligately cross-feeding co-culture is set by the MIC of the least tolerant species in the community. This phenomenon allows us to determine how antibiotics should interact in co-culture based on how they interact in each monoculture. A sample of these predictions are detailed in **Figure 2**. In brief, the co-culture is predicted to grow only where both species can grow individually (see plate diagrams). The impact of weakest link dynamics on antibiotic interactions depends on whether the weakest link species is the same or different in both antibiotics, and how the antibiotics interact with each species. In scenario 1, the weakest link species differs in each antibiotic, but in both species the antibiotic effects are independent of each other; therefore, the antibiotics should also be independent in co-culture. This is seen in the FICI plots (where the median FICI is around 1) and in the isobolograms (where the curve is around the 1-1 line). In scenario 2, the antibiotics synergize in both species, but because weakest link species differs in each antibiotic, the synergism is weakened (though still present) in co-culture. In scenario 3, the antibiotics antagonize in both species. However, in *E. coli*, antibiotic B antagonizes antibiotic A (i.e. as the concentration of B increases, the MIC of A also increases), but not vice versa (i.e. the MIC of B does not change as the concentration of A increases). In *S. enterica*, antibiotic A antagonizes antibiotic B but not vice versa. This leads to a ‘cancelling out’ of the antagonistic interactions in co-culture and causes the antibiotics to interact independently. In scenario 4, *E. coli* is the weakest link species in both antibiotics. Therefore, the co-culture antibiotic interaction pattern exactly matches that of *E. coli*.

**Figure 2.**
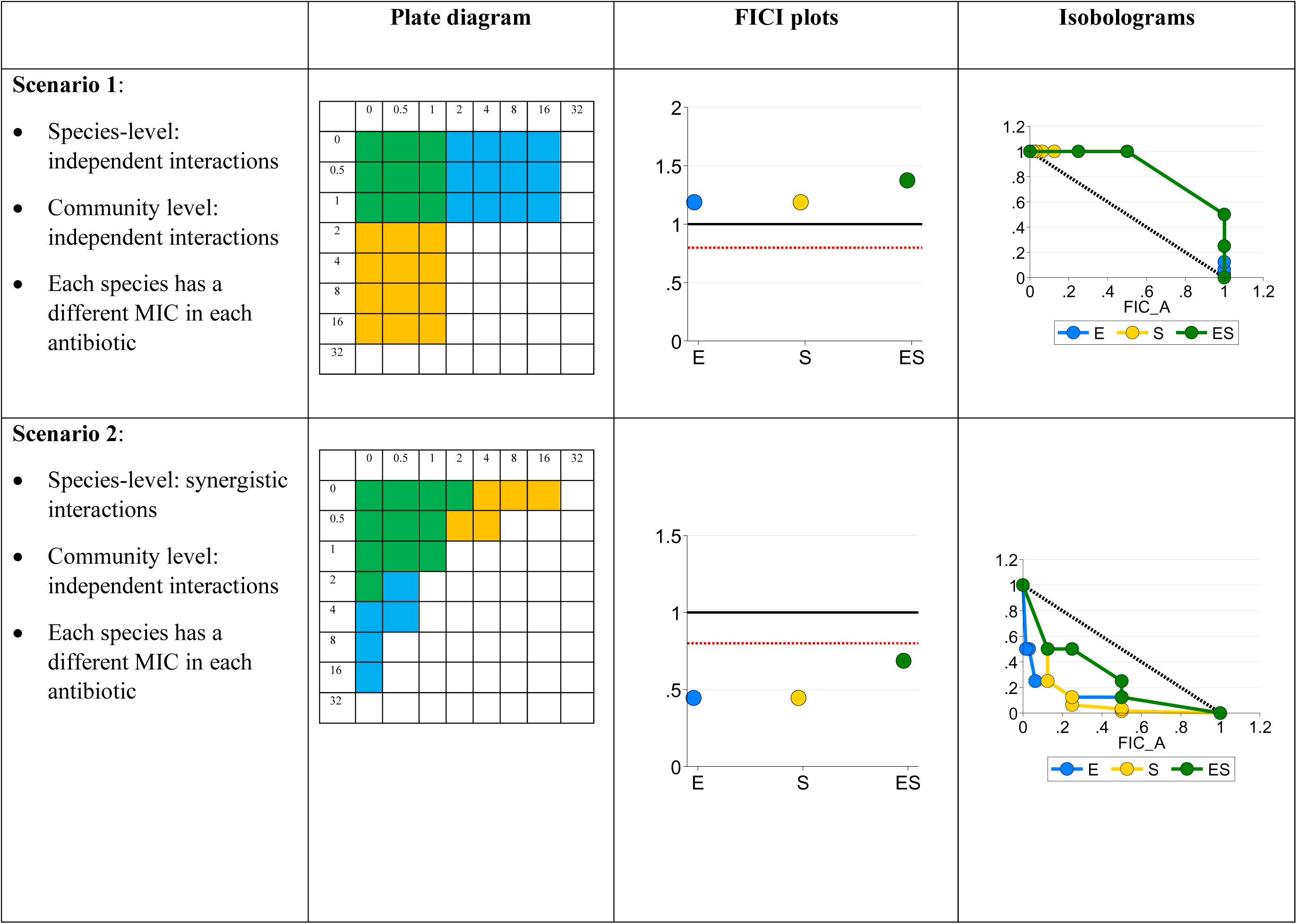
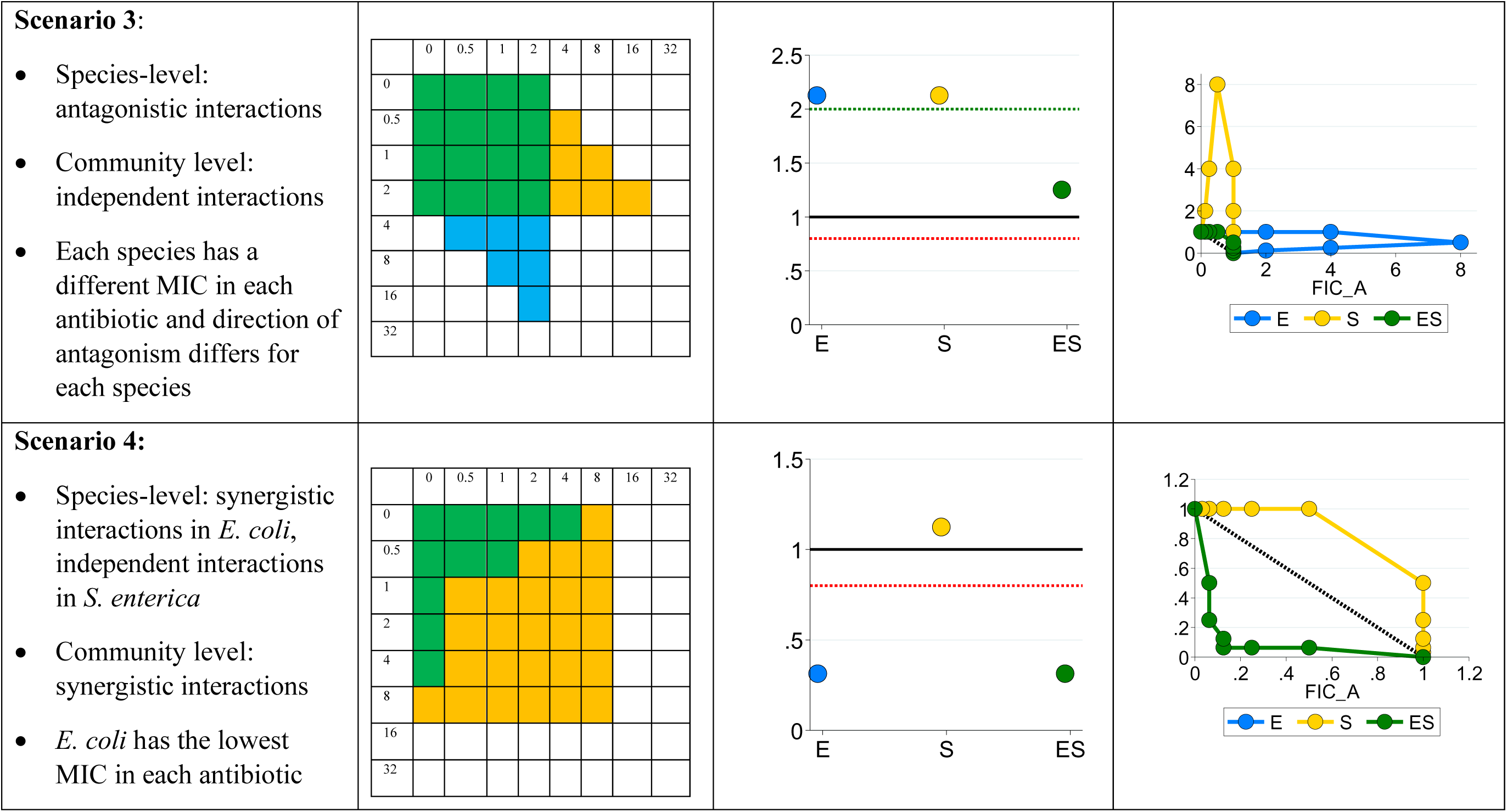
Antibiotic interactions at the species level versus the co-culture level. In the plate diagrams (simulated data), blue cells represent concentrations where only *E. coli* can grow; yellow cells represent concentrations where only *S. enterica* can grow, and green cells represent concentrations where the co-culture can grow (i.e. concentrations where both monocultures can grow). Antibiotic A is on the y-axis and antibiotic B is on the X-axis. Points that fall below the red dotted line on FICI plots represent synergistic interactions; points that fall above the green dotted line represent antagonistic interactions. FICI plots and isobolograms were calculated based on the simulated data in plate diagrams (see **Methods**). Concave isoboles represent synergy; convex isoboles represent antagonism.

We first tested whether the antibiotic combinations we selected would interact as predicted in the literature in our monocultures. We tested each antibiotic combination in triplicate for *E. coli* and *S. enterica*, then calculated the median FICI value for each plate and combination (**Figure 3**). Our categories were designated as follows: FICI < 0.8 represents synergy, FICI between 0.8 and 1 represent additive interactions, FICI between 1 and 2 represent independent interactions, and FICI ≥ 2 represents antagonism. These are less stringent than other FICI results because we chose median values to minimize the impact of plate-to-plate variation, and medians tend to bias FICI results away from detecting interactions. We also looked at isobolograms (**Figure 4**) of each antibiotic combination for each species, to get a more visual/qualitative examination of interactions between antibiotics. **Supplementary tables S2** and **S3** contain raw median and minimum FICI data, respectively.

**Figure 3.**
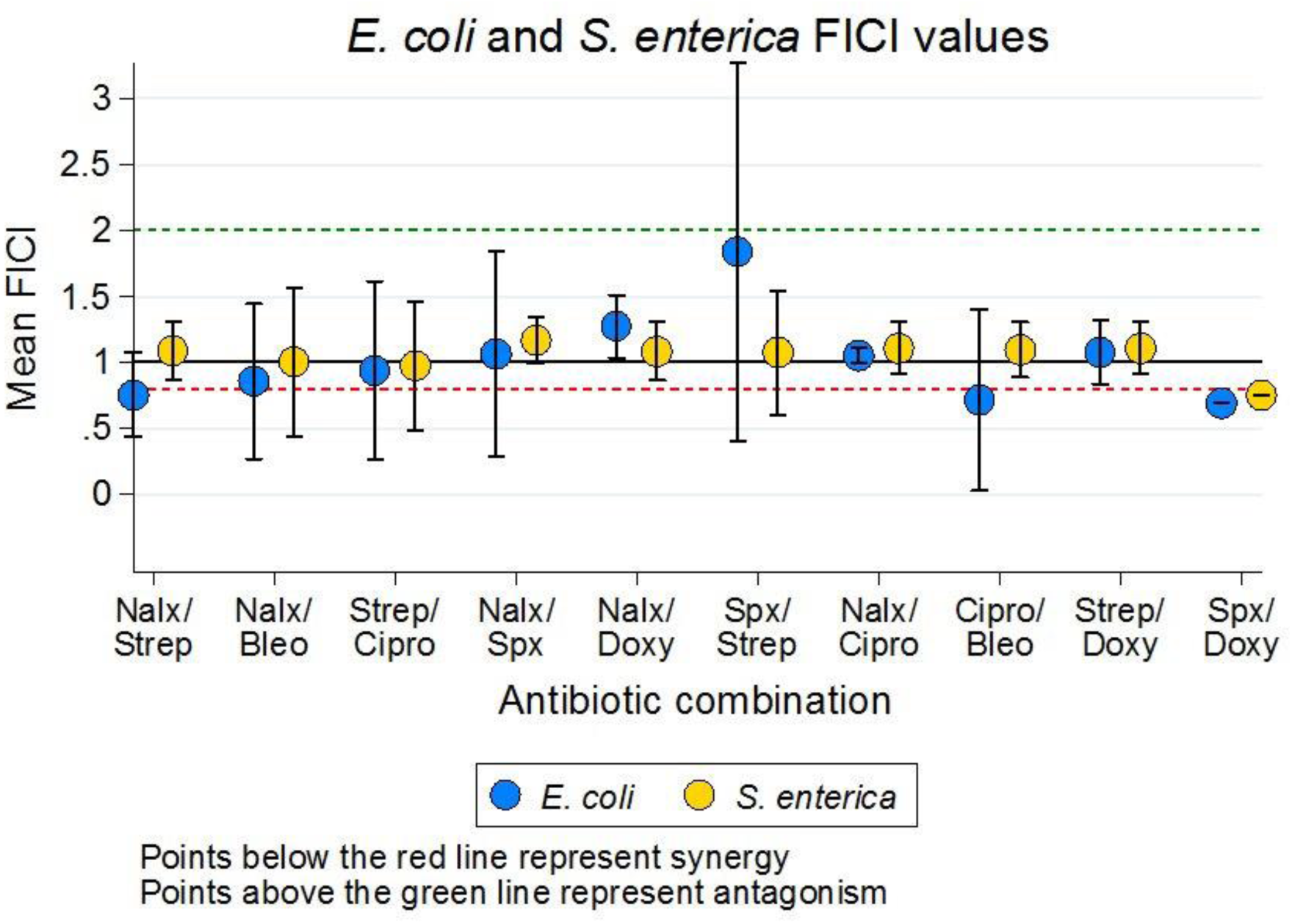
Fractional inhibitory concentration index (FICI) plots of *E. coli* and *S. enterica* monocultures across ten antibiotic combinations. Each point represents the mean +/-SE of three replicate FICI values from three biological replicates. FICIs on each plate represent the median FICI value from the plate. Antibiotic abbreviations: Nalx= nalidixic acid; strep= streptomycin; bleo= bleomycin; cipro= ciprofloxacin; spx= spectinomycin; doxy= doxycycline.

**Figure 4.**
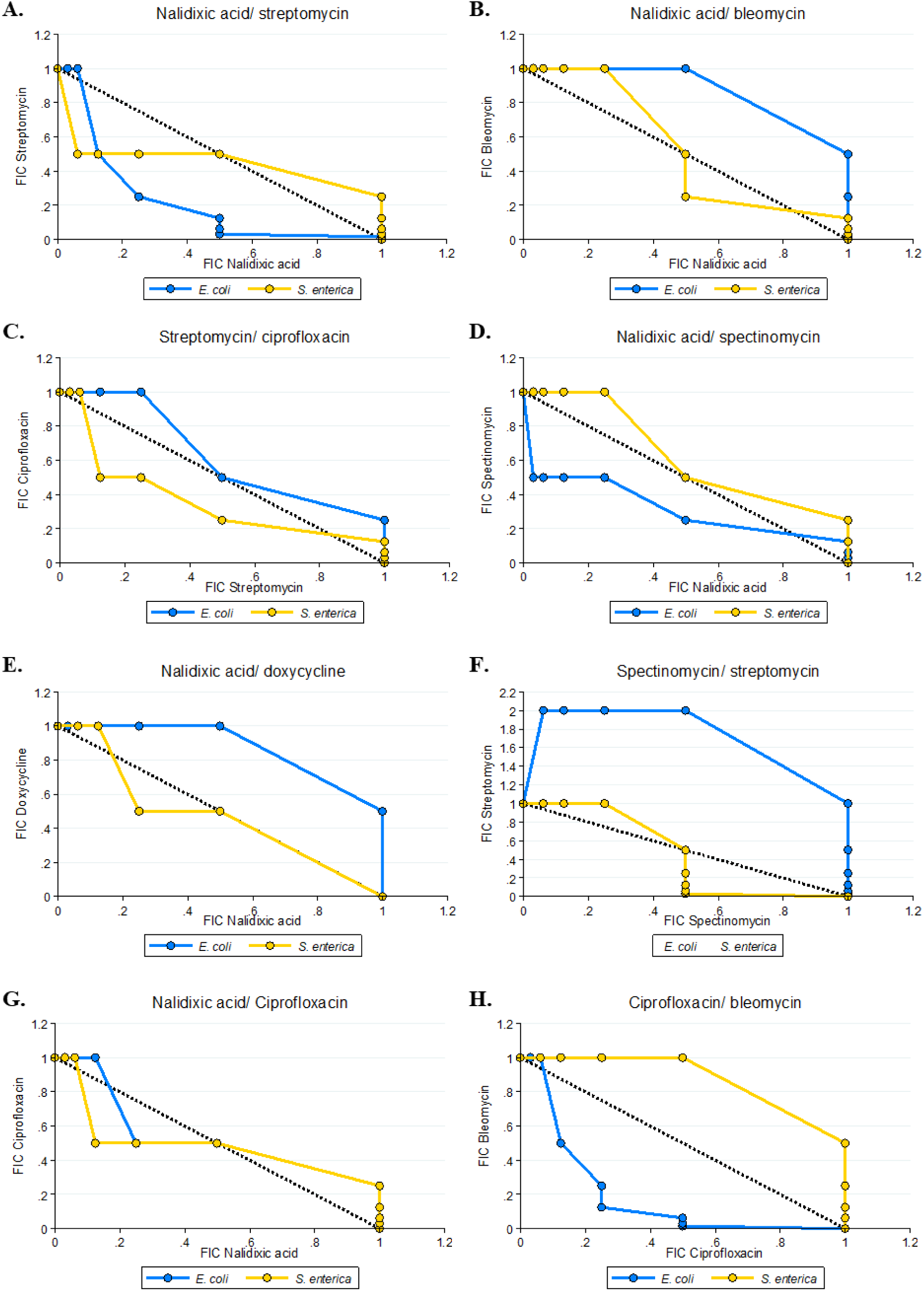
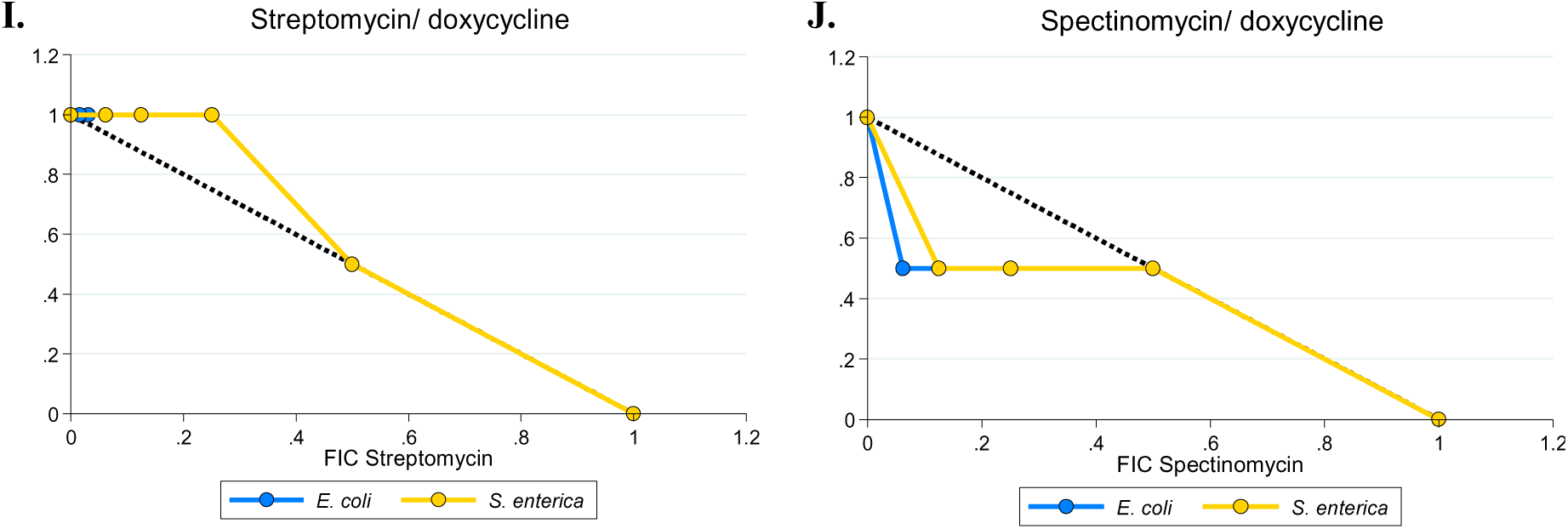
Representative isobolograms of *E. coli* and *S. enterica* monoculture fractional inhibitory concentrations (FICs) across ten antibiotic combinations. FICs were calculated based on 48 hours of 30°C growth, and growth was identified as any well which had an OD600 at least 10% of the highest OD600 well on each plate. Each axis corresponds to a fractional inhibitory concentration (FIC) for the antibiotic pair. The black 1-1 line represent a perfectly independent interaction; a concave line towards the origin represents a synergistic interaction, and a convex line away from the origin represents an antagonistic interaction.

Interestingly, we did see some deviations from our prediction. Nalidixic acid/bleomycin and streptomycin/ciprofloxacin were predicted to synergize; however, our FICI and isobologram data show additive/independent interactions for these antibiotics in both species. Nalidixic acid and streptomycin did synergize as predicted in *E. coli*, but not in *S. enterica*. Of the three pairs of antibiotics predicted to antagonize (nalidixic acid/ spectinomycin, nalidixic acid/doxycycline, and spectinomycin/streptomycin), only the last showed potentially antagonistic interactions; the others all interacted independently. Finally, we observed some unexpected synergy in our antibiotic pairs which were predicted to interact additively/independently.

Ciprofloxacin/bleomycin synergized in *E. coli*, and spectinomycin/doxycycline synergized in both species; however, this is more evident in the FICI data than in the isobolograms. The isobolograms suggest that low concentrations of doxycycline decrease the MIC of spectinomycin, but not vice versa; that is, doxycycline synergizes with spectinomycin to increase the latter’s potency, but spectinomycin does not change the effect of doxycycline.

Based on our results from monoculture and our weakest link hypothesis, we then predicted the antibiotic interactions which would arise in obligate cross-feeding co-culture. To generate these predictions, we examined the monoculture growth patterns in each antibiotic combination (i.e. at which concentrations of each antibiotic monoculture growth occurred). We then generated a predicted growth pattern for the co-culture wherein growth would only occur at antibiotic concentrations where both species could grow. From this predicted growth pattern, we calculated FICIs and generated isobolograms; these can be seen in **Figure 5** and **6**, respectively. An example of how this was done can be found in **Supplementary figure S1.**

**Figure 5.**
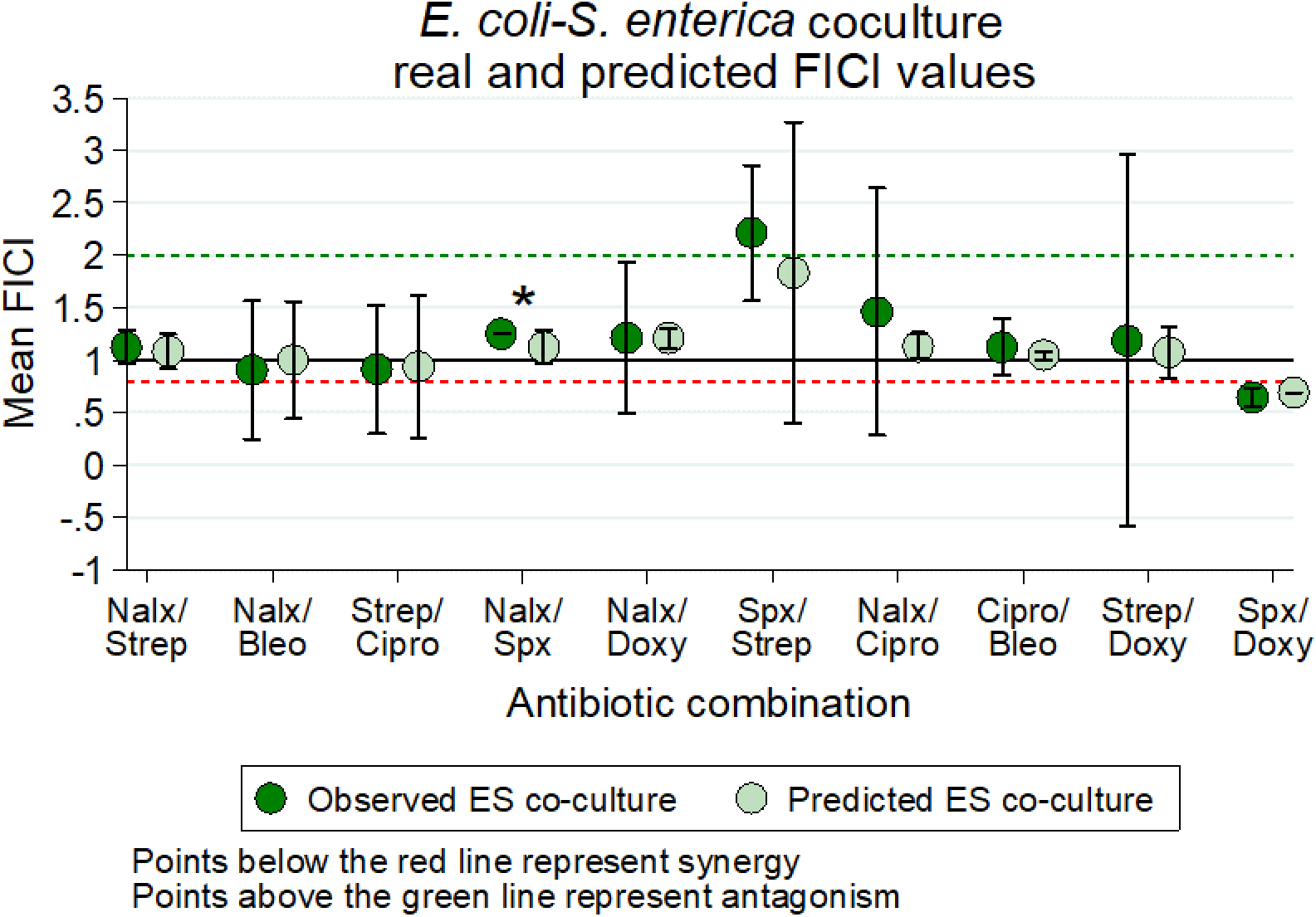
Fractional inhibitory concentration index (FICI) plots of predicted and actual co-cultures across ten antibiotic combinations. Each point represents the mean +/-SE of three replicate FICI values from three biological replicates. FICIs on each plate represent the median FICI value from the plate. Asterisks represent *P* < 0.05 for predicted versus observed ES co-culture FICs were compared with a Mann-Whitney U test. *P*-values can be found in **Supplementary table S6**. Antibiotic abbreviations: Nalx= nalidixic acid; strep= streptomycin; bleo= bleomycin; cipro= ciprofloxacin; spx= spectinomycin; doxy= doxycycline.

**Figure 6.**
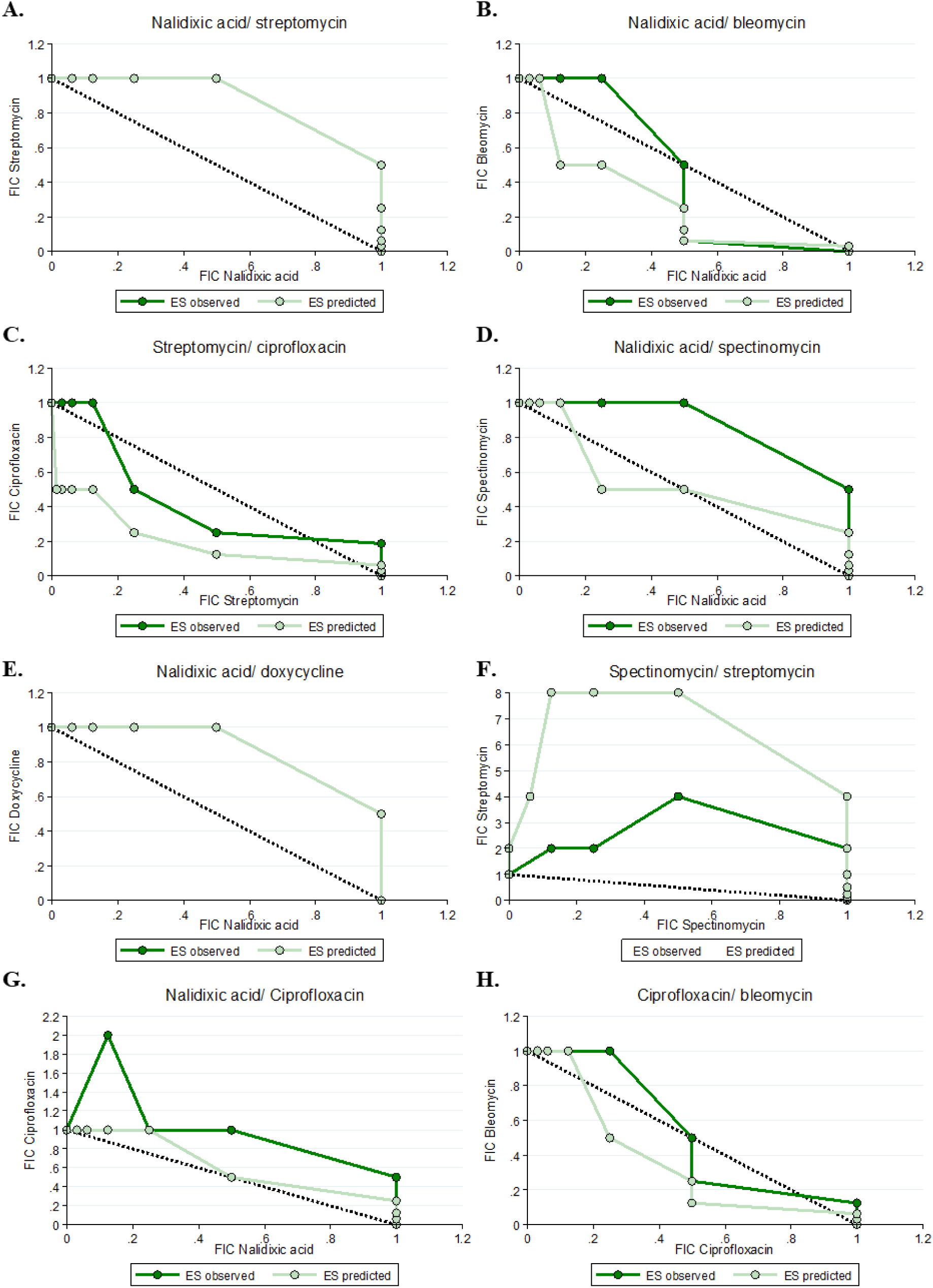
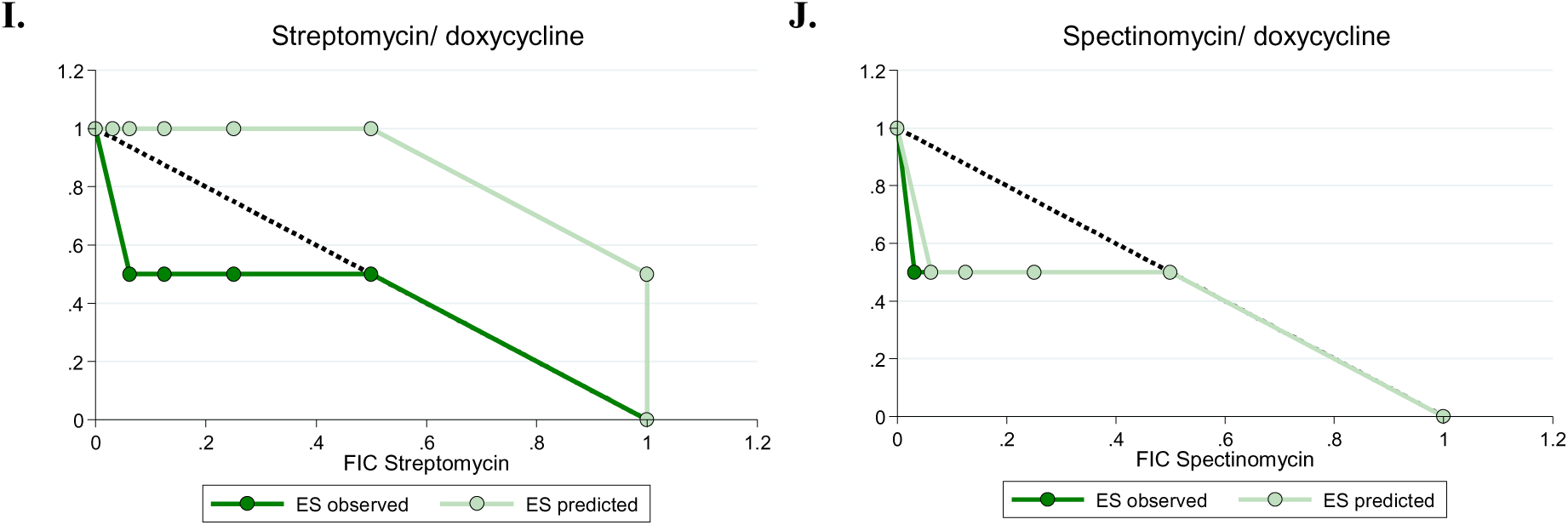
Representative isobolograms of predicted and observed co-culture fractional inhibitory concentrations (FICs) across ten antibiotic combinations. Predicted FICs were determined based on monoculture FICs and hypothesized weakest link dynamics (i.e. co-culture growth could only occur at concentrations of both antibiotics where both species could grow alone). Observed co-culture FICs were calculated based on 48 hours of 30°C growth, and growth was identified as any well which had an OD600 at least 10% of the highest OD600 well on each plate.

According to our predictions, if one species is the weakest link (i.e. the least tolerant) in both antibiotics, the co-culture interaction typically matched that of the weakest link monoculture. This is the case for nalidixic acid/bleomycin and nalidixic acid/ciprofloxacin (where *S. enterica* is the weakest link), and for streptomycin/ciprofloxacin, spectinomycin/streptomycin, streptomycin/doxycycline, and spectinomycin/doxycycline (where *E. coli* is the weakest link). Co-culture predictions were somewhat more complicated for the other combinations (nalidixic acid/ streptomycin, nalidixic acid/spectinomycin, nalidixic acid/doxycycline, and ciprofloxacin/ bleomycin), where each species is the weakest link in a different antibiotic. We were particularly interested in nalidixic acid/streptomycin, as these antibiotics synergize in *E. coli* (which is the weakest link in streptomycin) and interact independently in *S. enterica* (which is the weakest link in nalidixic acid). Based on the differences in MIC in these species in each antibiotic (see **Supplementary table S4**), we predicted an independent interaction in co-culture. Similarly, in the ciprofloxacin/ bleomycin combination, the antibiotics verged on antagonizing in *E. coli* and interacted independently in *S. enterica*; however, their MICs were similar in both antibiotics. This provided an opportunity to examine interactions in co-culture where weakest link dynamics might play less of a role.

After generating predicted FICIs based on our monoculture results and weakest link dynamics, we tested antibiotic interactions in co-culture. We then compared our predicted FICIs to those observed experimentally for each antibiotic combination. Qualitatively, our predictions based on weakest link were accurate — the antibiotic interaction category (antagonism/synergy/additive) identified by predicted FICIs matched the interaction category identified by the observed FICIs (**Figure 5,** see **Supplementary Table S5** for raw FICI data). This supports our hypothesis that weakest-link dynamics can be used to predict antibiotic interaction categories in co-culture. The one exception to this was in the spectinomycin/ streptomycin combination. While there was no statistical significance in this difference, (*P*= 0.37), we predicted an independent interaction and observed an antagonistic interaction. Interestingly, the isobologram suggested that antibiotics antagonized much more in co-culture than we predicted. This suggests that weakest-link dynamics may not always predict co-culture outcomes and that some other factor may be determining antibiotic interactions in this case. Quantitatively, our FICI predictions also matched that of our observed data (see **Supplementary Table S6** for all *P-*values), with one exception. The predicted FICI for the nalidixic acid/ spectinomycin combination was significantly higher than predicted (*P*= 0.037), but this difference still resulted in independent interactions and so is likely not biologically significant. Overall, weakest-link dynamics were generally sufficient to both qualitatively and quantitatively predict antibiotic interactions in co-cultures.

## Discussion

The goal of this work was to identify whether our previously identified weakest link hypothesis, wherein the antibiotic tolerance of a mutualistic co-culture is set by the weakest link species, could change drug interaction patterns in antibiotic combinations. We tested previously identified antibiotic combinations in each of our monocultures. Few of the predicted interactions applied to our monocultures, possibly for reasons discussed below. However, we then used the interactions we identified in monoculture, as well as our knowledge of weakest link dynamics, to predict how each set of antibiotics would interact in co-culture. We found that our predictions were qualitatively correct, with predicted and observed FICIs and isobolograms falling into the same antibiotic interaction category (synergistic, additive, independent, or antagonistic). The one exception to this was the spectinomycin/streptomycin combination, which antagonized more strongly in co-culture than we predicted from monoculture.

Our findings demonstrate an important and hitherto unexplored explanation for why *in vivo* antibiotic interactions do not match *in vitro* assay predictions. Many infections are now known to be polymicrobial (27, 28, 35, 39) and likely involve some form of cooperative metabolite exchange. These ecological interactions may be at least partially responsible for the difficulty in finding a successful synergistic antibiotic treatment. Indeed, our results suggest that cross-feeding generally ablates any antagonistic/ synergistic antibiotic interactions unless one partner is the weakest link in every antibiotic (**Figure 2**); whether or not this is the case in natural microbial communities is unknown. Helpfully, our results suggest that the antibiotic interactions at the community level are predictable given the right information — i.e. if the individual resistances and antibiotic interaction patterns are known for each species in the community, the antibiotic interaction pattern is generally predictable based on weakest link dynamics. This adds further weight to the argument that microbial ecology must be considered when treating bacterial infections in the clinic.

Unexpectedly, the antibiotic interactions that we observed in our monocultures did not match the interactions that Yeh et al. had previously observed (16). The most likely reason for this is their use of a growth rate-based measurement method, a dose-response curve (12), versus our yield-based checkerboard assay. We elected to do a yield-based method because it allowed us to more highly parallelize our experiments and decrease plate-to-plate variation in cell density and growth phase, both of which are known to significantly impact antibiotic tolerance (40–42). Much research has been done on the best method for assessing antibiotic synergy/antagonism (12, 43, 44); we selected the checkerboard method also because of its widespread use and ease of interpretation. Future experiments using dose-response curves might be particularly important for cross-feeding systems such as ours, as cross-feeding is known to alter growth rates of member species (45, 46).

An additional challenge in interpreting antibiotic interactions in multispecies contexts is the possibility of antibiotic interactions changing depending on which species is the weakest link at a given combination of antibiotic combinations. Taking the largest FICI value from a plate biases results towards antagonism and taking the smallest value biases towards synergy. Therefore, the median value is useful in avoiding overinterpretation of data; however, it obscures any concentration-specific changes in interactions which might be occurring. We reported isobolograms and FICIs for this reason. Isobolograms provide more information as to how the antibiotics are interacting at different concentration combinations than FICIs. The isobologram of nalidixic acid/bleomycin in **Figure 6** provides a good example of this. The predicted co-culture isobole showed additive-synergistic interactions; however, the observed co-culture isobole showed synergistic interactions at low bleomycin FIC values. A similar pattern is seen with ciprofloxacin/bleomycin in the same figure. While these patterns may be artifacts of our system, it remains possible that checkerboard assays involving multiple species may produce isobologram patterns which deviate from the typical convex/concave, antagonism/synergy pattern seen in monocultures. Mathematical modeling of how different antibiotic interactions and MICs in each species impact co-culture antibiotic interactions may be a useful way to explore this possibility.

The one drug interaction in our study where weakest link dynamics appeared insufficient to predict co-culture interactions was the streptomycin/spectinomycin combination. These drugs were predicted to antagonize in *E. coli*; though they have similar mechanisms of action, spectinomycin ionically inhibits entry of streptomycin into the cell (47). Given that *E. coli* was the weakest link in both antibiotics, we predicted similar dynamics in co-cultures; additive interactions bordering on antagonism (i.e. FICIs between 1.5 and 2). However, the degree of antagonism that we observed was much higher than predicted. There could be several reasons for this. Given that disruptions in protein biosynthesis have pleiotropic effects on cell physiology and metabolism (48), the application of both drugs might have sufficiently disrupted the cross-feeding between our species such that they starved at otherwise sublethal concentrations of each antibiotic. That antibiotics can arrest growth rate (49, 50) and change metabolic profile (51, 52) of cells is well known; what is less clear is how this might impact metabolite exchange in antibiotic-exposed natural microbial communities. The complex and often non-obligate metabolite exchange food webs in natural communities (53, 54) might make this question difficult to answer, but our study suggests that weakest link dynamics are a useful null hypothesis starting point.

Though much research has been done *in vitro* on antibiotic synergy/antagonism, it remains unclear what the biological/clinical relevance of any of these interactions truly are. With a few exceptions (2, 7), antibiotic synergy has yet to be adopted as a clinically important treatment strategy despite some success in mouse models (55, 56). Differences in drug half-life and bioavailability can impact effective dosages *in vivo* (57), and strain-specific resistance profiles make assessment of antibiotic synergy challenging in the clinic (12). However, antibiotic combinations may become a critical clinical tool as resistance continues to rise (13). Further research is therefore required not just on how antibiotics interact *in vitro*, but how they interact in natural environments— both within the host, and within a multispecies community.

## Methods

Our model microbial community has been previously described (37). Briefly, our system consists of an *E. coli* methionine auxotroph, and an *S. enterica* strain which has been evolved to secrete excess methionine. In a lactose environment, *E. coli* metabolizes lactose to produce acetate for *S. enterica*, which in turn supplies methionine for *E. coli*. Each species can also be grown in monoculture by supplying *E. coli* with methionine and lactose, and *S. enterica* with acetate.

We performed checkerboard assays (described below) with six antibiotics in ten different combinations predicted to synergize (3), antagonize (3), or not interact (4)— see **Table 1** for these combinations. For each drug combination, we tested *E. coli* and *S. enterica* in monocultures, and the two-species in obligate co-culture. Each antibiotic combination/culture type was tested in triplicate. Seven two-fold dilutions of each antibiotic, along with an antibiotic– free control for each, were used in orthogonal gradients on a 96-well plate such that the antibiotic concentrations increased from left-to-right and top-to-bottom. The first row and column of each plate were antibiotic–free wells for the vertically– and horizontally– distributed antibiotics, respectively. The minimum inhibitory concentrations (MICs) for each antibiotic were determined in the absence of the other antibiotic. Mid-log–phase cells (OD∼0.4) were grown up on the day of the experiment in species–specific Hypho growth medium (36) and 2µL was inoculated into 194µL fresh species-specific Hypho. Antibiotic stocks were prepared within two days of the experiment such that 2µL of stock could be added to each well to achieve the desired gradient concentrations. Plates were then incubated at 30°C with shaking for 48 hours. A Tecan plate reader was then used to measure the OD600 and species-specific fluorescence (CFP for *E. coli* and YFP for *S. enterica*). The 90% minimum inhibitory concentration (MIC_90_) was then used to establish which wells showed growth. Any well that had an OD600 or fluorescent protein value above 10% of the highest plate value was considered growth. We used the highest plate value rather than the antibiotic–free well because we consistently saw a slight increase in OD600 in the co-cultures at sublethal concentrations, possibly due to a low level of cell lysis and subsequent boost for the cross-feeding partner (58, 59).

We used the Loewe additivity method to identify the nature of our antibiotic interactions as previously described (6). Briefly, we calculated the fractional inhibitory concentration (FIC) for antibiotics A and B as follows: FIC_A_ = (MIC_A in combination_ / MIC_A alone_), and FIC_B_ = (MIC_B in combination_ / MIC_B alone_). FIC values were obtained for each well at the edge of growth, as shown in **Figure 1.** The FICI is the sum of FIC_A_ and FIC_B_ (60). As there are multiple FICI values per plate, we chose to report the median FICI value as the plate value. We did not use the minimum or maximum FICI value so that we would not over-interpret synergy or antagonism results, respectively (61). Minimum FICI values can be found in **Supplementary table S3**. Our cut-off values were designed as follows: FICI < 0.8 represents synergy; FICI between 0.8 and 2 represent additive interactions, FICI between 1 and 2 represent independent interactions, and FICI ≥ 2 represents antagonism (60–63). Isobolograms were generated by plotting the FIC_A_ and FIC_B_ values as x,y coordinates. A straight line connecting the FIC values represents additive interactions; a concave line represents synergy; and a convex line represents antagonism.

Based on observed monoculture growth patterns (MICs and FICs in each antibiotic combination), we predicted co-culture growth patterns assuming weakest link dynamics; that is, co-cultures should only grow at concentrations of both antibiotics where both species are able to grow in monoculture. We then calculated FICs and FICIs for these predicted co-culture plates and compared them to our observed data. We then used a Mann-Whitney U test to compare predicted versus observed FICIs for our co-cultures.

## Acknowledgements

The authors would like to thank Jeremy Chacón, Lisa Fazzino, Sarah Hammarlund, Brian Smith, and Leno Bernard Smith Jr. for their insights on this work. This work was supported by a Natural Sciences and Engineering Research Council of Canada Postgraduate Scholarship (PGSD2-487305-2016, to E.M.A) and by a National Institutes of Health Award (1R01-GM121498, to W.R.H.).

## Figures and Tables

**Supplementary table S1.** Mechanism of action of antibiotics used in this study.

| Antibiotic | Mechanism |
| --- | --- |
| Bleomycin | Induces DNA breaks; may inhibit thymidine incorporation into DNA |
| Ciprofloxacin | Fluoroquinolone: binds DNA gyrase and topoisomerase IV |
| Nalidixic acid | Naphthyridone: binds DNA gyrase and topoisomerase IV |
| Doxycycline | Binds 30s ribosomal subunit to prevent protein biosynthesis |
| Spectinomycin | Binds 30s ribosomal subunit to prevent protein biosynthesis |
| Streptomycin | Binds 30s ribosomal subunit to prevent protein biosynthesis |

**Supplementary table S2.** Median FICIs for *E. coli* and *S. enterica* in monoculture across ten antibiotic combinations and three replicates. FICIs for each replicate are the median FICI value per plate. FICI values below 0.8 are considered synergy; FICIs between 0.8 and 1 are additive interactions, FICIs between 1 and 2 are independent interactions, and FICIs above 2 are antagonistic interactions.

| Species | Antibiotic combination | Rep 1 | Rep 2 | Rep 3 |
| --- | --- | --- | --- | --- |
| *E. coli* | Nalidixic acid/ streptomycin | 0.75 | 0.625 | 0.8828125 |
|  | Nalidixic acid/ bleomycin | 0.6875 | 0.75 | 1.125 |
|  | Streptomycin/ ciprofloxacin | 1.125 | 0.625 | 1.0625 |
|  | Nalidixic acid/ spectinomycin | 1.0625 | 0.75 | 1.375 |
|  | Nalidixic acid/ doxycycline | 1.25 | 1.1875 | 1.375 |
|  | Spectinomycin/ streptomycin | 2.5 | 1.5 | 1.5 |
|  | Nalidixic acid/ ciprofloxacin | 1.0625 | 1.0234375 | 1.0625 |
|  | Ciprofloxacin/ bleomycin | 0.546875 | 0.5625 | 1.03125 |
|  | Streptomycin/ doxycycline | 1 | 1.046875 | 1.1875 |
|  | Spectinomycin/ doxycycline | 0.6875 | 0.6875 | 0.6875 |
| *S. enterica* | Nalidixic acid/ streptomycin | 1.1875 | 1.015625 | 1.0625 |
|  | Nalidixic acid/ bleomycin | 1.1875 | 0.75 | 1.0625 |
|  | Streptomycin/ ciprofloxacin | 1.125 | 0.75 | 1.03125 |
|  | Nalidixic acid/ spectinomycin | 1.25 | 1.125 | 1.125 |
|  | Nalidixic acid/ doxycycline | 1.03125 | 1.03125 | 1.1875 |
|  | Spectinomycin/ streptomycin | 1.25 | 0.875 | 1.09375 |
|  | Nalidixic acid/ ciprofloxacin | 1.1875 | 1.09375 | 1.03125 |
|  | Ciprofloxacin/ bleomycin | 1.1875 | 1.03125 | 1.0625 |
|  | Streptomycin/ doxycycline | 1.1875 | 1.09375 | 1.03125 |
|  | Spectinomycin/ doxycycline | 0.75 | 0.75 | 0.75 |

**Supplementary table S3.**
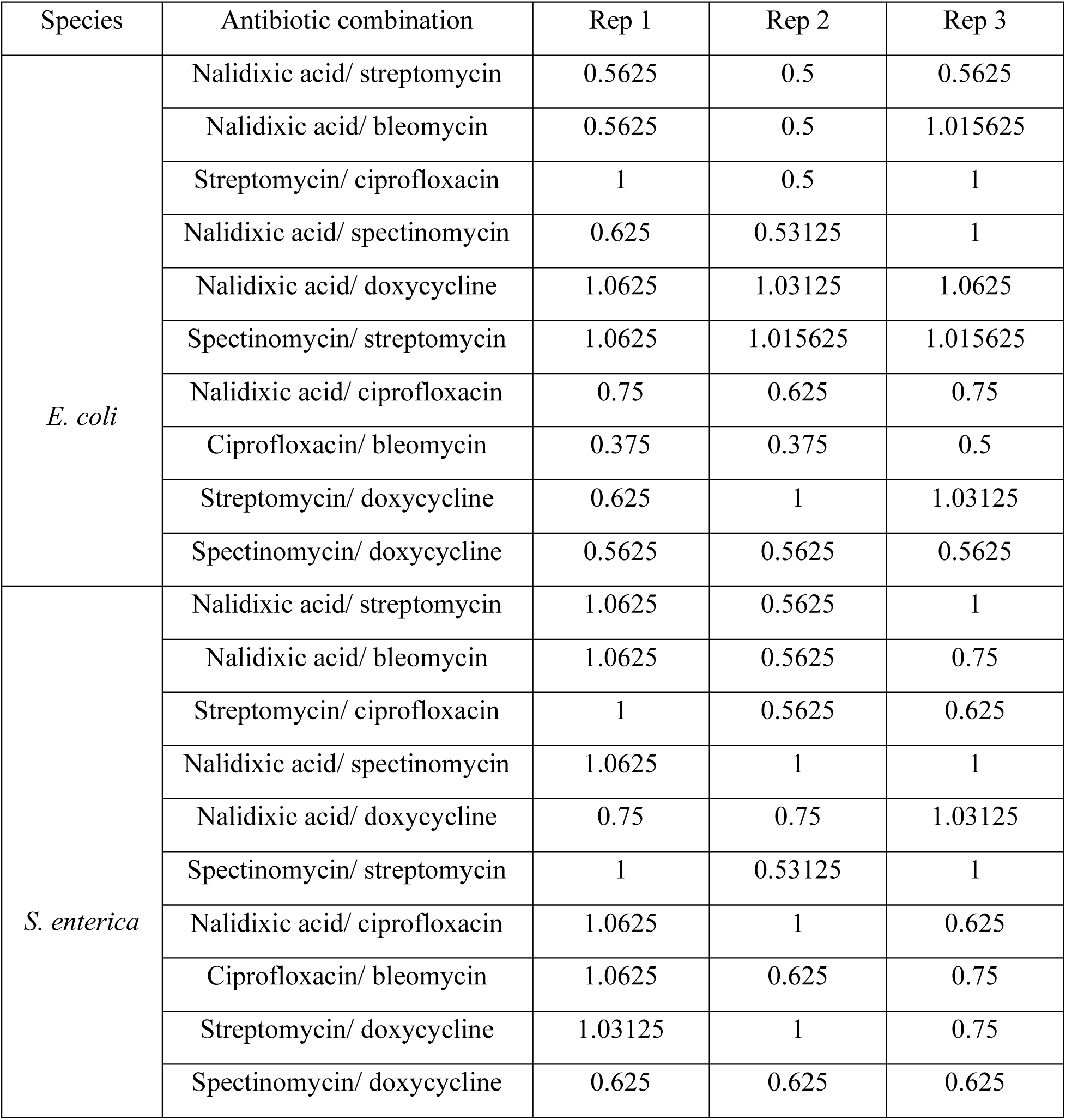
Minimum FICIs for *E. coli* and *S. enterica* in monoculture across ten antibiotic combinations and three replicates. FICIs for each replicate are the minimum FICI value per plate. FICI values below 0.8 are considered synergy; FICIs between 0.5 and 1 are additive interactions, FICIs between 1 and 2 are independent interactions, and FICIs above 2 are antagonistic interactions.

**Supplementary figure S1.**
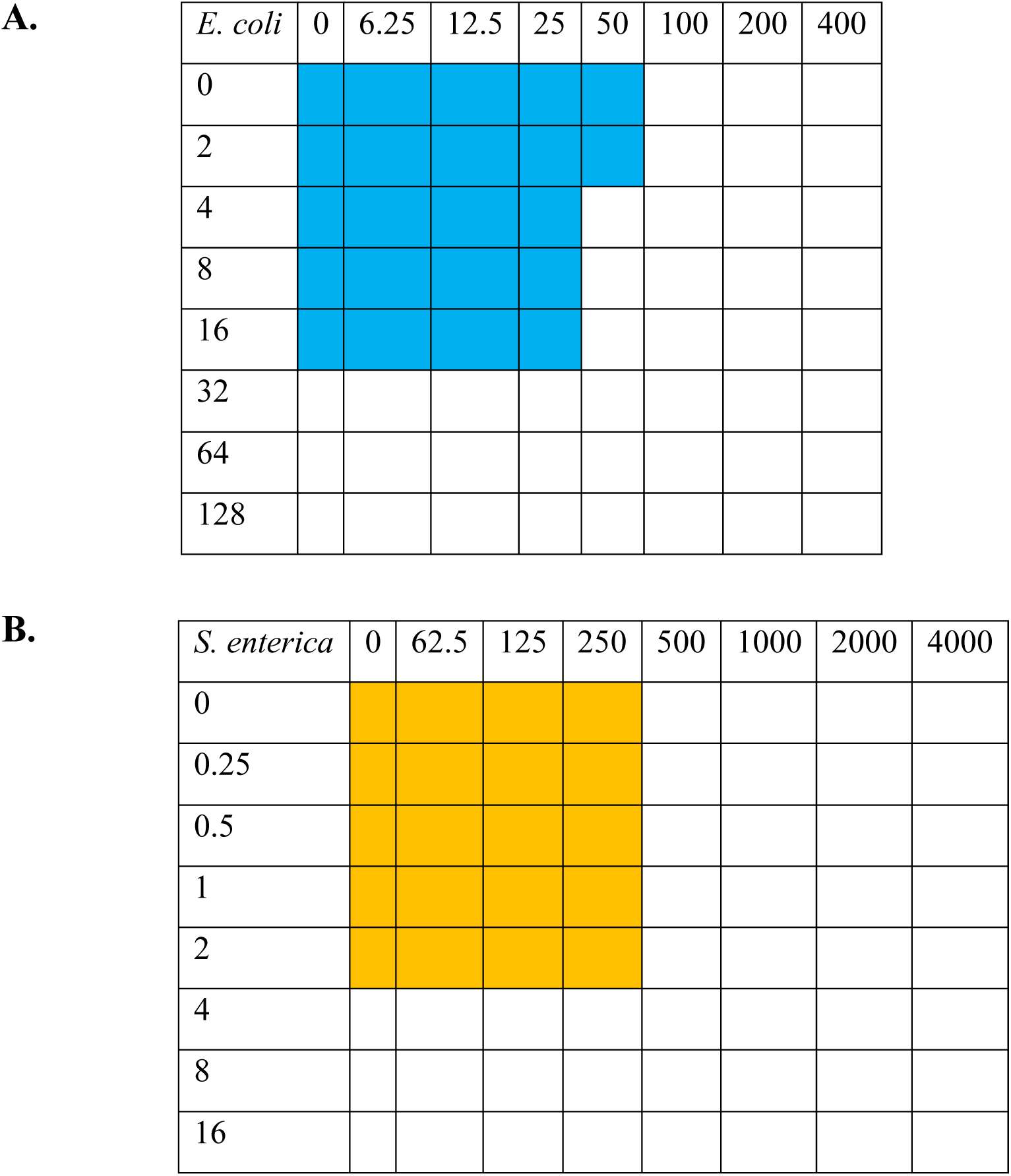
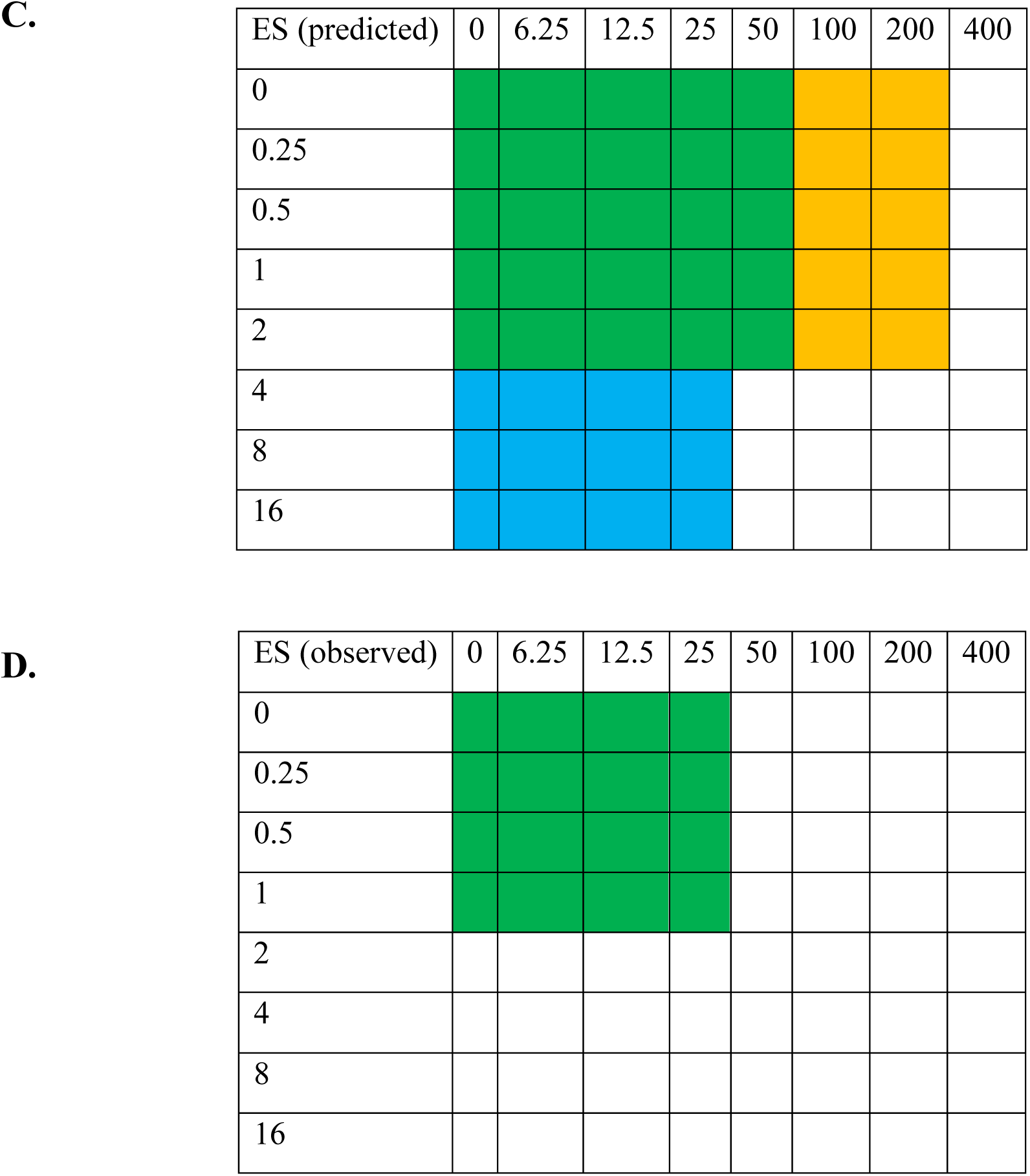
Example of developing predicted FICIs from replicate 1 of nalidixic acid/ spectinomycin combination. Growth patterns of *E. coli* (**A**) and *S. enterica* (**B**) monocultures were used to predict growth patterns for the co-culture (**C**). FICIs and isobolograms were developed from this predicted data as previously described, and these were compared to real data obtained from co-cultures (**D**).

**Supplementary table S4.4.** Minimum inhibitory concentrations (MICs) of each species in each antibiotic, predictions for co-cultures based on weakest link, and actual co-culture MICs. MICs were defined as the lowest concentration of antibiotic required to inhibit growth below 10% of the densest well (by OD600) within a plate. Medians and ranges are displayed. Predicted co-culture MICs are based on weakest link hypothesis (i.e. the co-culture will be limited by the least resistant monoculture).

| **Antibiotic** | ***E. coli* MIC** | ***S. enterica* MIC** | **Predicted co-culture MIC** | **Observed co-culture MIC** |
| --- | --- | --- | --- | --- |
| Bleomycin (µg/mL) | 8 (2-8) | 2 (1-2) | 2 (1-2) | 1 (0.5-2) |
| Ciprofloxacin (ng/mL) | 16 (8-32) | 16 (16-32) | 16 (8-32) | 16 (8-16) |
| Doxycycline (µg/mL) | 0.25 (0.0625-0.25) | 2.5 (2.5-5) | 0.25 (0.0625-0.25) | 0.25 (0.125-0.25) |
| Nalidixic acid (µg/mL) | 32 (32-64) | 8 (4-8) | 8 (4-8) | 2 (1-4) |
| Spectinomycin (µg/mL) | 100 (100-200) | 500 (500-1000) | 100 (100-200) | 100 (50-200) |
| Streptomycin (µg/mL) | 1.5 (0.5-2) | 160 (80-160) | 1.5 (0.5-2) | 0.5 (0.5-8) |

**Supplementary table S5.**
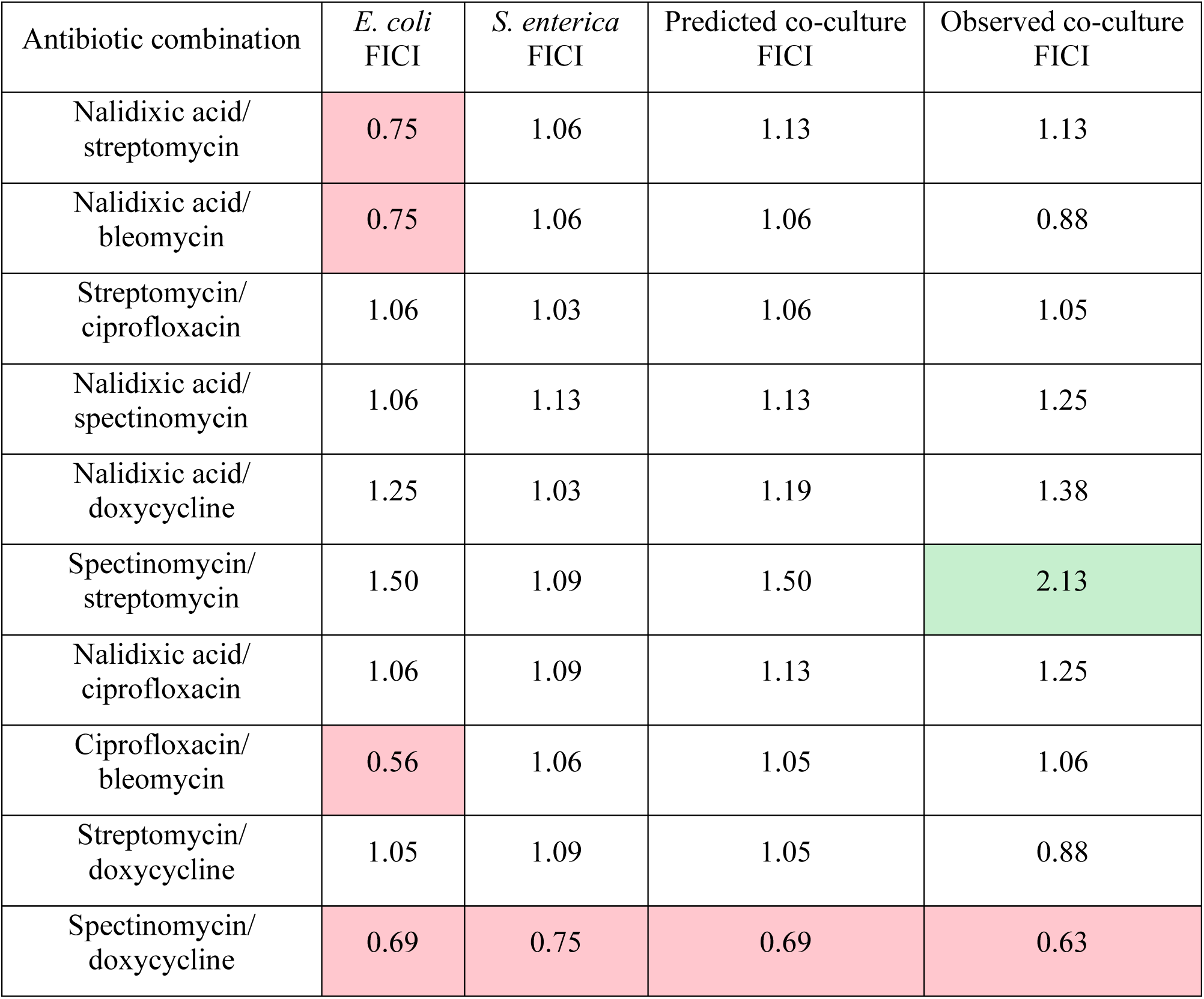
Observed fractional inhibitory concentration indices (FICIs) for each antibiotic combination in monoculture and co-culture, and predicted co-culture FICIs based on weakest link. FICIs are median values from three biological replicates each. Red cells represent synergistic interactions (median FICI<0.8); green cells represent antagonistic interactions (median FICI>2).

**Supplementary table S6.** Mann-Whitney U statistical test results for predicted vs. observed FICI results.

| Antibiotic combination | *P*-value for predicted vs. observed FICI |
| --- | --- |
| Nalidixic acid/ streptomycin | 0.49 |
| Nalidixic acid/ bleomycin | 0.66 |
| Streptomycin/ ciprofloxacin | 0.50 |
| Nalidixic acid/ spectinomycin | 0.037 |
| Nalidixic acid/ doxycycline | 0.50 |
| Spectinomycin/ streptomycin | 0.37 |
| Nalidixic acid/ ciprofloxacin | 0.18 |
| Ciprofloxacin/ bleomycin | 0.10 |
| Streptomycin/ doxycycline | 0.51 |
| Spectinomycin/ doxycycline | 0.11 |

